# Towards 3D-bioprinting of osseous tissue of pre-defined shape using single-matrix cell-bioink constructs

**DOI:** 10.1101/2022.07.27.501781

**Authors:** Yawei Gu, Sebastian Pigeot, Lucas Ahrens, Fabian Tribukait-Riemenschneider, Melika Sarem, Francine Wolf, Andrea Barbero, Ivan Martin, V. Prasad Shastri

## Abstract

Engineering living bone tissue of defined shape on-demand has remained a challenge. 3D-bioprinting (3DBP), a biofabrication process capable of yielding cell constructs of defined shape, when combined with developmental engineering can provide a possible path forward. Through the development of a bioink possessing appropriate rheological properties to carry a high cell load and concurrently yield physically stable structures, printing of stable, cell-laden, single-matrix constructs of anatomical shapes was realized without the need for fugitive or support phases. Using this bioink system, constructs of hypertrophic cartilage of predesigned geometry were engineered in vitro by printing human MSCs at a high density to drive spontaneous condensation and implanted in nude mice to evoke endochondral ossification. The implanted constructs retained their prescribed shape over a 12-week period and underwent remodeling to yield ossicles of the designed shape with neovascularization. Micro-CT, histological and immunohistochemistry assessments confirmed bone tissue characteristics and the presence of human cells. These results demonstrate the potential of 3DBP to fabricate complex bone tissue for clinical application.

## Introduction

Bone grafts are required to reconstruct bone defects caused by trauma, ablative surgeries, and congenital abnormalities. Human bones can be classified into five categories based on their shapes, which are related to their distinct functions ^1^. Beside an obvious mechanical function, bones in the maxillofacial region also serve an aesthetic purpose and therefore impact patients’ mental health ^2,3^. Thus, engineering bone tissue with anatomical features is pivotal in restoring holistic attributes following bone loss. Autologous bone graft, which is the gold standard, is associated with increasing donor-site morbidity and a limited source ^4^. Although strategies such as the cell-free “in vivo” bioreactor, or cell-scaffold based tissue engineering through intramembranous ossification (IO) can replicate bone tissues, these approaches are not well-suited to engineer living bones of precise shape ^5,6^. Since, IO requires active recruitment of neo-vasculature, bone formation through endochondral ossification (EO) is more attractive, as it starts with a cartilaginous template that is inherently avascular, and therefore can survive implantation and since the provisional hypertrophic cartilage matrix is pro-angiogenic it can undergo remodeling into bone ^7–9^. 3D bioprinting (3DBP) with EO has been explored using MSCs printed in gelatin methacrylate (GelMA) and alginate, and in these studies bone tissues were found in the printed constructs after in vivo implantation ^10,11^. However, both systems require secondary processing, i.e., photo crosslinking in the case of GelMA to stabilize the print, or a polycaprolactone internal physical scaffold to support the weak ionically crosslinked alginate phase. The need for secondary processing increases the complexity of the biofabrication process itself. An alternative approach would be to employ bioinks that can yield self-supporting structures thereby giving access to single-matrix constructs.

Previously, we have demonstrated that bioinks formulated with carboxylated-agarose (CA) ^12^ could be used to print free-standing complex structures without fugitive phases or post-processing ^13^. Additionally, it has been shown in a recent study that CA hydrogels support neo-vascularization and stabilization of blood vessels ^14^. Based on these encouraging results here we explored if CA-based bioinks can be used to pre-define the shape of the bone tissues formed through EO. However, in order to exploit 3DBP using CA bioinks two criteria need to be met: 1) the printed structures should incorporate high-density of cells without impairing the printability, since in engineering tissues, a high volumetric density of cells is a prerequisite to ensure homogenous tissue deposition and for activating cellular processes dependent on cell-cell contact such as tubulogenesis, epidermal sheet formation, MSC condensation ^15,16^; and 2) the printed MSC-CA structures should be converted into hypertrophic cartilage templates for the EO program to be invoked in vivo. Towards this objective, in this study CA formulation that supports printing high density of cells was identified and explored as a bioink for engineering bone tissue through EO (**Figure 1**).

**Fig. 1.**
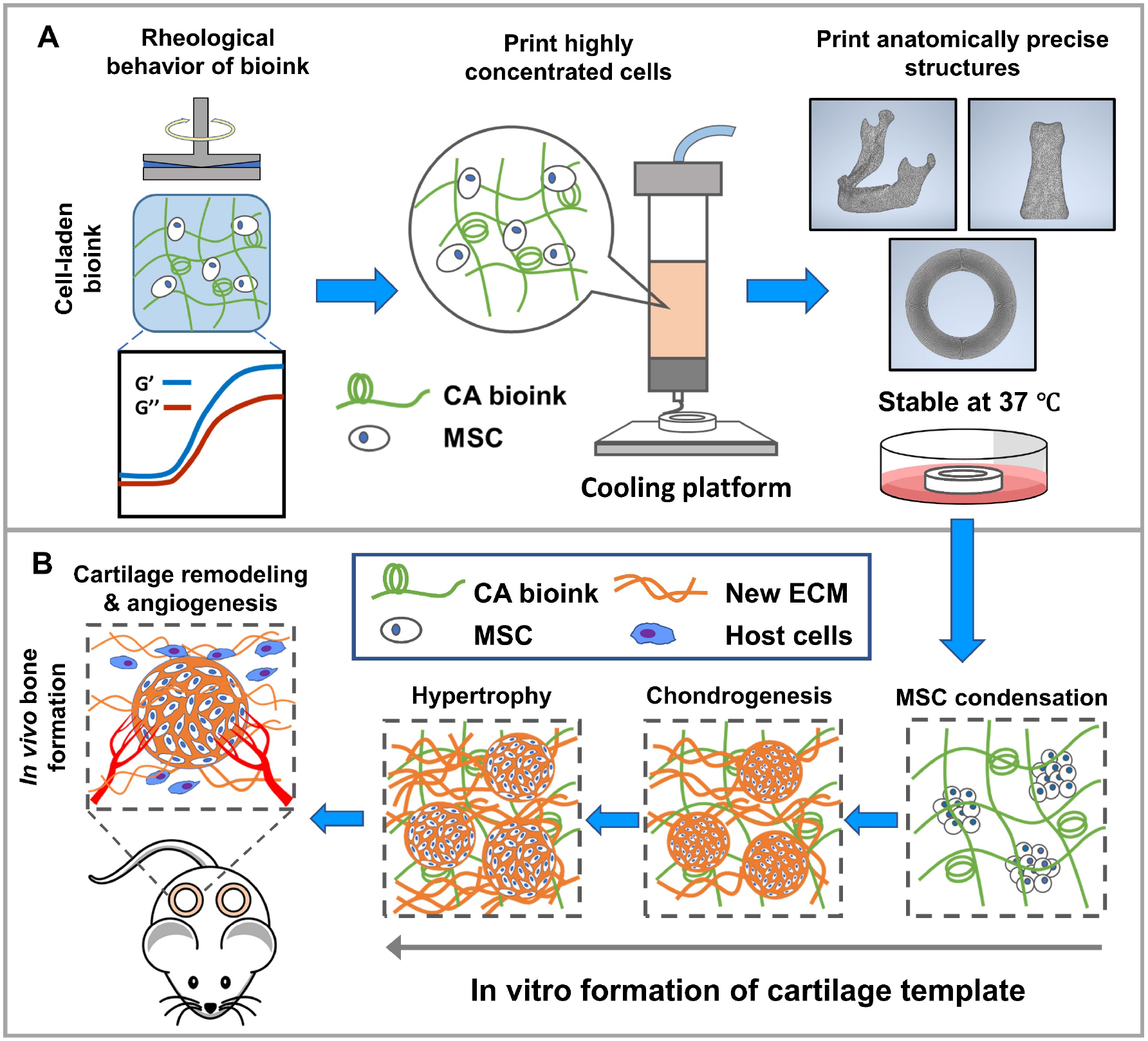
Study Design. **(A)** Identification through rigorous rheological testing of a bioink formulation that is capable of carrying a high concentration of cell and is suitable for printing constructs in shapes consistent with bone structures from various anatomical locations without support or fugitive materials, or post-processing. Finite element mesh representation of a human mandibular bone, a human 3^rd^ middle finger phalange and a ring are shown as exemplar structures for bioprinting. **(B)** De novo aggregation of human MSCs printed within 3D-bioprinted ring-shaped constructs and differentiation into hypertrophic cartilaginous template, followed by implantation in athymic mouse for remodeling through endochondral ossification (EO) to bone in the shape defined *a priori*.

## Results

### Carboxylated agarose (CA) bioinks support high concentration of cells without losing printability

Building on our recent study showing that CA-hydrogel based bioink can yield free-standing structures such as bifurcated tubes or hemispheres without the need for a support or fugitive phase, and post-processing such as chemical crosslinking ^13^, we embarked on optimizing this system to make it suitable for carrying a high density of cells towards the development of a single-matrix construct for bone engineering. In our print workflow, the bioink is first equilibrated at 45 °C before mixing with the cells, transferred to a pre-heated cartridge to 37 °C, and then printed onto a print bed cooled to 4 °C, following which the printed structure is transferred to a 37 °C incubator for culturing (**Fig. 2A**). It has been shown that the volumetric contribution of a large cell number is not negligible, and can influence the rheological behavior of polymer-cell solutions, as well as impair crosslinking ^17^. In the CA system, introducing a high concentration of cells could influence the gelation behavior by impeding formation of physical crosslinks. We undertook a detailed characterization of the bioink in absence of cells (cell-free) and in presence of cells (cell-laden, 60 × 10^6^ cells/mL), keeping in mind that such a comparative analysis would be also valuable for standardization toward clinical translation ^18^, and ascertained the influence of cells on the storage (G’) and loss (G”) modulus of cell-laden and cell-free bioink using a series of rheological tests to simulate the entire fabrication process (bioink preparation and printing). We identified that a composite of two biopolymers, CA and native agarose at 19:1 weight ratio, when formulated at 10 w/v% yielded a bioink system that showed no appreciable changes in both G’, G” at both 45 °C, at 37 °C and retained its gelation behavior even upon addition of cells (**Fig. 2B**). Frequency sweep tests at the printing temperature (37 °C) revealed that the addition of cells transformed the cell-laden bioink into a more viscous composite (**Fig. 2C**), while still retaining gel-like properties of the cell-free bioink as confirmed by an elastic modulus that was higher than loss modulus. Although holding the bioink at 37 °C for 30-minute led to an increase of viscosity in both cell and cell-free bioinks, despite higher viscosity, the yield stress of cell-laden bioink was lower than cell-free bioink, which is consistent with behavior reported in collagen bioink ^19^, and this can be translated into lower extrusion pressure to print cell-laden bioink, which is an important attribute of the CA bioink system (**Fig. 2D, 2E**). As a consequence, cells experience less induced-shear stress during printing ^12^.

**Fig. 2.**
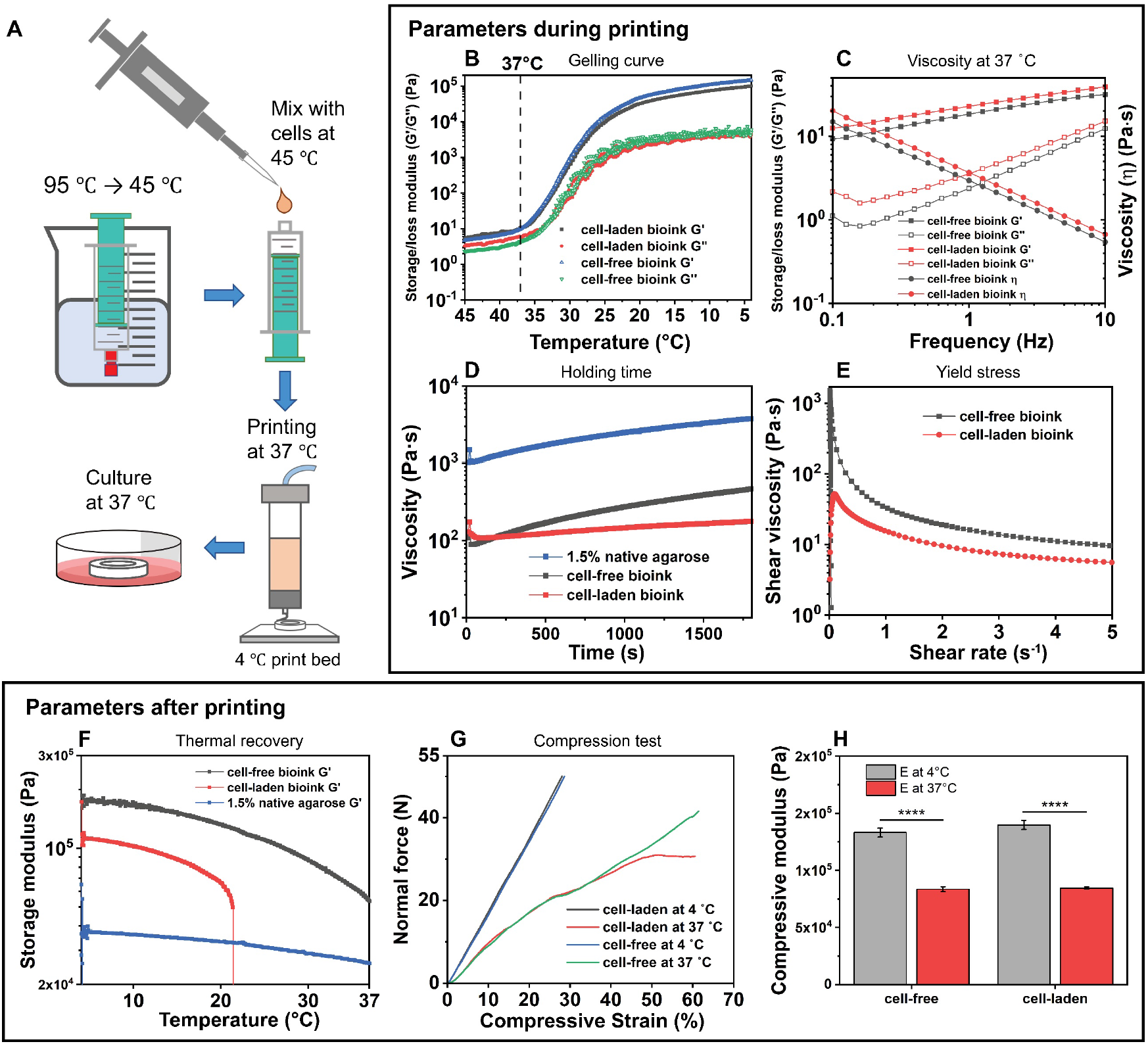
Advanced CA-based bioink is capable of printing cells at high densities without impairing printability and mechanical stability. (A) The work flow of preparation of carboxylated agarose (CA) bioink and bioprinting. Upper box: Rheological properties relevant during printing. (B) Gelling behavior of bioinks during a temperature sweep ranging from 45 °C to 4 °C. (C) The viscosity of bioinks at printing temperature (37 °C) when oscillation exerted at frequency from 0.1 Hz to 10 Hz. (D) Influence of holding time in cartridge on the viscosity of bioinks. I Yield stress of bioinks at 37 °C. Lower box: Rheological and mechanical properties relevant post-printed structures. (F) Thermal recovery behavior of bioinks during a temperature sweep range from 4 °C to 37 °C, which simulated the transfer from print bed to cell incubator. (G) Stress-strain (compressive) curves of bioinks at 4 °C and 37 °C. (H) Compressive modulus of bioink at 4 °C and 37 °C. Notes: The curves of 1.5 w/v% native agarose in C and E are set as reference.

In addition to rheological properties, the ability to implant 3D-bioprinted structures hinges on the structural stability of printed structures. In this regard, one consideration is the thermal relaxation in CA-hydrogels leading to softening of the hydrogel, due to the weak hydrogen bonding between β-sheet structures in CA backbones ^**13**^. While the presence of cells led to a sharp drop in storage modulus around ambient temperature, plausibly due to a plasticization effect imparted by the cells promoting softening of gels (**Fig. 2F**), this did not compromise the integrity of the gel as evidenced next. The compressive moduli at 4 °C and at 37 °C of both non-cell laden bioink (133.1 ± 3.8 kPa and 83.4 ± 2.2 kPa) and cell-laden bioink were similar (139.7 ± 3.9 kPa and 84.4 ± 1.1 kPa), suggesting that the bioink formulation was well suited for printing stable cell laden structures (**Fig. 2G, 2H**).

### Printing anatomically accurate structures without support phases and post processing

The CA-based bioink was explored for its utility to reproducibly yield structures ranging from simple 2D patterns to complex 3D structures including scaled-down models of human-skeletal anatomy. Two basic 2D structures with biological relevance such as of an array of hexagons (segment length = 2.2 mm) and a sheet composed of 2-3 layers of several closely spaced lines were printed, confirming the ability of subsequent layers to fuse with one another (**Fig. 3A-D**). Since structures such as bone in extremities are not uniformly or symmetrically shaped form the distal-to-proximal end, we next explored the printing of high aspect ratio structures, for example a tube (30 mm high) starting and ending with two equilateral triangles that are rotated by 60° rotation with respect to each other, resulting in a twisted motif. To increase the complexity further and mimic dimensional transitions in long bone, a spherical region was incorporated in the middle of the print (**Fig. 3E, F**). When properly rendered, such a structure presents a star-of-David configuration when viewed from the top and a tortuous body when viewed from the side (**Fig. S2A, B**). Encouraged by the successful printing of this complex object, the printing of objects that are anatomically accurate with respect to their dimensional relationship and shape was undertaken. Towards this objective three structures were chosen: the human lumbar vertebral body (**Fig. 3G, H**), the mandibular bone (**Fig. 3I, J**) and the middle phalange of the third finger (**Fig. 3K, L**).

**Fig. 3.**
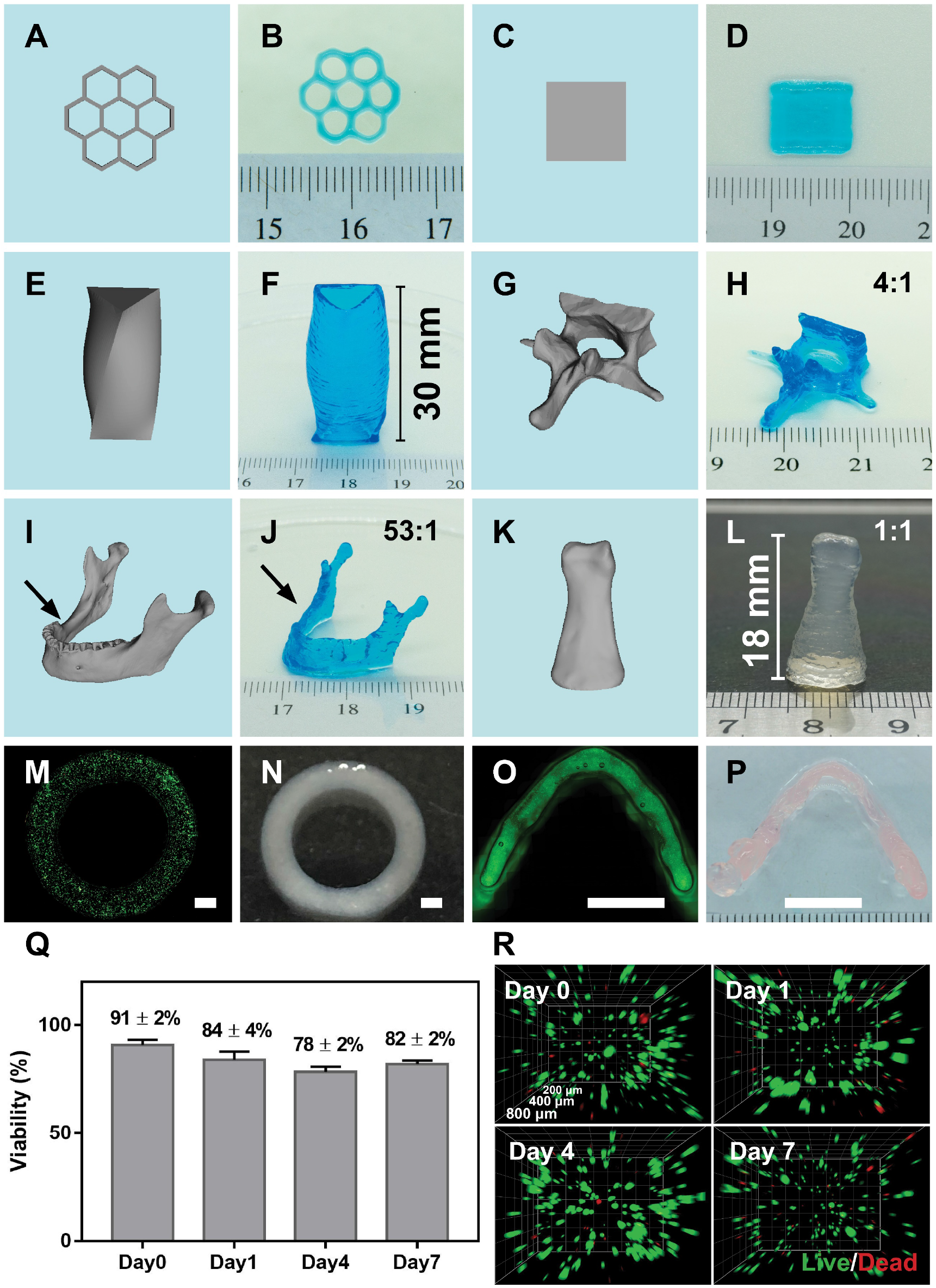
3D bioprinted structures possess high fidelity and resolution, while affording high cellular viability. The 3D models of all the printed structures are presented in A, C, E, G, I, and K. CA-based bioink is capable of printing 2D structures like a hexagon-assembly and a square sheet with high resolution (B, D), an irregular tube with two triangular openings rotated by 60° with respect each other (F shows the side view), scale-down anatomy-precise structures like the first lumbar vertebral body (H) and a mandibular bone (J), and the 3^rd^ middle phalange in human scale (18 mm × 9 mm × 6.5 mm) (L). Black arrows in I and J show the bone defect in the position that spanned the second premolar to third molar. The precision of ruler in all figures is millimeter (mm). Cells distributed homogeneously inside printed structures with high viability after printing (M, O), and the printed constructs maintained stable during culture (N, P). O and P show the bottom view and top view of a scale-down construct of mandibular bone, respectively. Viability of MSCs after printing from day 0 to day 7 (Q). Representative 3D reconstructed fluorescent image of printed samples from day 0 to day 7, with green color showing viable cells and red color showing dead cells. Cell concentration in the bioink was 1 × 10^6^ cells/mL. The ratio in H, J and L is the ratio between actual size in human and the size of printed constructs. Scale bars in M, N are 1 mm, and scale bars in O, P are 10 mm. (n = 3 in Q and R)

The choice of the structures was based on the consideration that while the former two involve complex transitions in shape, thickness and orientation of the bone are generally high aspect ratio objects, the latter is a solid structure. Furthermore, the mandibular bone had a bone defect in the position that spanned the second premolar to third molar. Scaled-down anatomical replicates of the vertebral body (4:1) and the mandibular bone (53:1) and a “**to scale**” (1:1) replica of the phalange were all successfully printed, thus exalting the utility of the bioink system for “bone on demand” 3D-printed human engineered tissues.

### CA bioink permits aggregation and chondrogenic differentiation of MSCs

Cell distribution inside the constructures, maintenance of cell viability, and stability are the fundamental requirements for a bioink to be suitable for tissue engineering and transplantation. As shown in Fig. 3M and 3O, cells (stained by Calcein-AM) distributed homogeneously inside the bioink, and the constructs maintained stable during culture (**Fig. 3N, P**). The MSC viability within printed structures (1 × 10^6^ cells/mL of bioink), determined qualitatively using live/dead staining, was over 90% immediately after printing and settled at around 80% the following days (**Fig. 3Q, R**).

MSC aggregation and condensation represent the first step in chondrogenesis and bone formation via EO, and it has been shown that MSC chondrogenesis is highly dependent on cell numbers within these aggregates ^20^. One strategy is to use micro-well culture to precisely pre-form MSC aggregates of defined cell number ^21^ and then introduce the aggregates into the bioink. Such an approach presents many practical challenges including time consuming and labor-intensive harvesting of the aggregates, potential disintegration of the aggregates during the introduction into the bioink and during the printing and need to have high volumetric density of aggregates to achieve homogenous distribution in a printed structure. Differing from past studies ^11^, here a bioink without any cell adhesion promoters such as RGD was used, to encourage MSC-MSC interaction and promote MSC aggregation. MSC expressing tdTomato fluorescent protein were seeded at densities of 10 × 10^6^, 30 × 10^6^, and 60 × 10^6^ cells per mL. Immediately following introduction into the hydrogel bioink (Day 0), MSC were distributed homogeneously and presented a round morphology (**Fig. S1A**). Day 3 onwards, cells shrank and assumed a smaller round shape (**Fig. S1A**), and on day 7 obvious cell aggregates could be found inside the hydrogel (**Fig. S1A, Fig. 4A**). Moreover, as cell density increased, the fluorescence showed a progressive shift to higher intensity values with broadening of the peak, implying that cells were more clustered at higher densities. (**Fig. S1B, Fig. 4B**). While as expected, on the day of seeding, the ratio of cell aggregates increased with cell density (**Fig. 4C, Fig. S3**), over time although the ratio of cell aggregates in the 10 × 10^6^/mL group remained unchanged beyond day 3, increasing aggregation was observed in 30 × 10^6^/mL and 60 × 10^6^/mL groups (**Fig. 4C, Fig. S3**), and cell aggregates in the 60 × 10^6^/mL group were larger and showed a wider distribution in aggregate size in comparison to the 30 × 10^6^/mL group (**Fig. 4D**). To identify a suitable cell density for MSC-chondrogenesis in 3DBP structures, MSCs were printed at 30 × 10^6^/mL and 60 × 10^6^/mL concentration within round discs (6mm diameter × 3mm high), differentiated under chondrogenic conditions and compared with MSCs-seeded in Ultrafoam™ collagen scaffold (6mm diameter × 3mm high round disc), an established reference material from previous studies ^20,22,23^. After 3 weeks of chondrogenic differentiation, both concentrations led to comparable glycosaminoglycan (GAG) deposition although less than in Ultrafoam™ (**Fig. 4E**). With further two weeks of hypertrophic culture, ECM distributed was more homogenous in the 3D bioprinted samples, and since the higher concentration of 60 × 10^6^/mL did not show clear benefits, the concentration of 30 × 10^6^/mL, which was also reported by Scotti et al., ^22^ was chosen for the bioprinting.

**Fig. 4.**
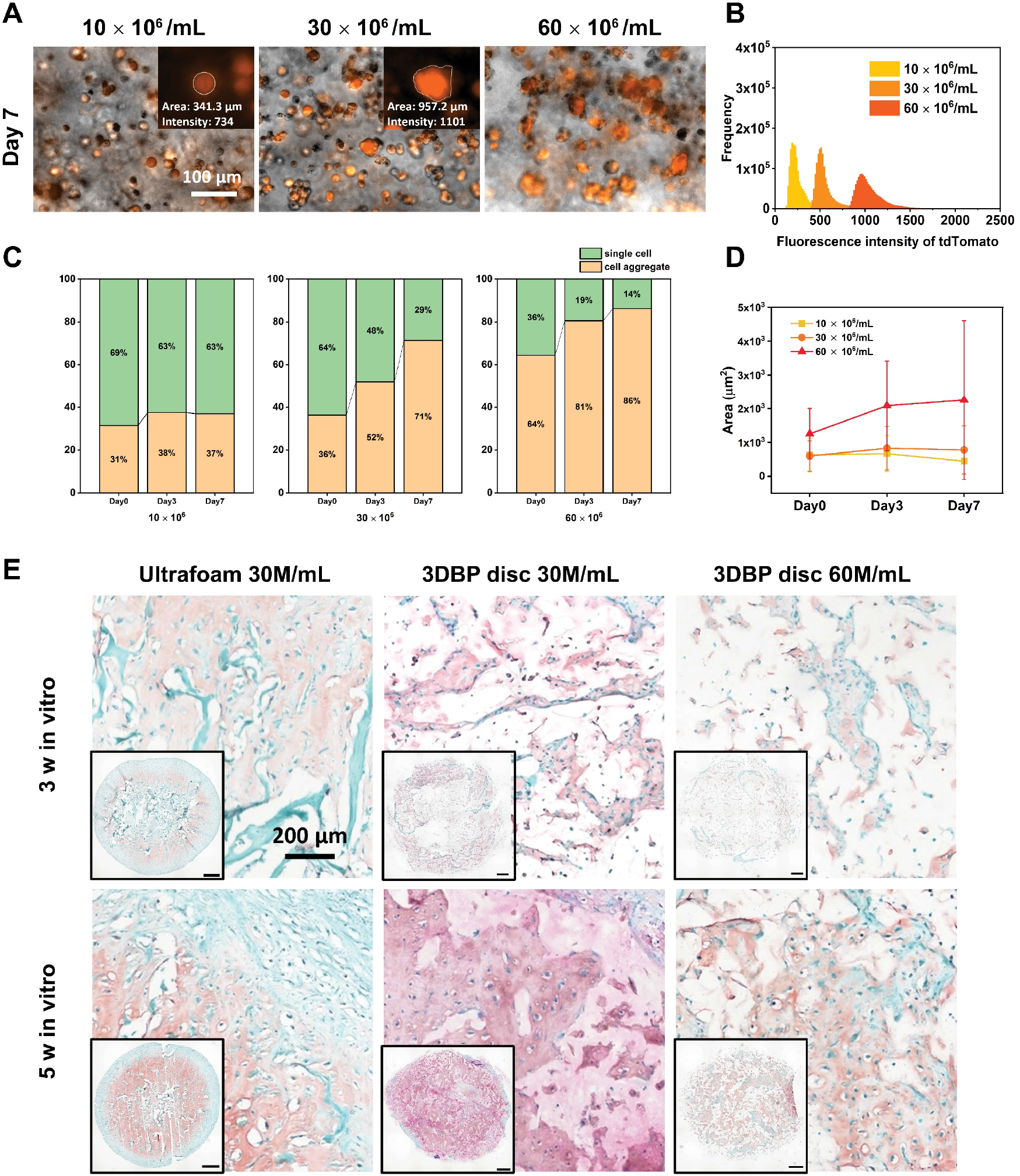
CA-bioink promotes spontaneous aggregation of human MSCs and supports chondrogenic differentiation. (A) Images with merged brightfield and fluorescence channels showing aggregation of MSCs at various densities within CA bioink after 7 days, with the insets showing the patterns of a single cell and cell aggregates. (B) Histogram showing distribution of fluorescence intensity of MSCs within the bioink on day 7 (X-axis fluorescence intensity, Y-axis frequency). (C) The percentage of cell aggregates and single cell in samples with different MSC density as function of culture time. (D) Change in average size of cell aggregates normalized to area of a single cell at different cell density as a function of culture time. (E) Safranin-O staining for sulfated glycosaminoglycans (s-GAGs) within Ultrafoam and 3D bioprinted discs with different concentrations of MSCs at the end of the chondrogenic (3 weeks) or the hypertrophic culture (5 weeks). Scale bar = 1 mm. (n = 3 in A-D, n = 3 in E)

### Formation of hypertrophic cartilage tissue by MSCs in 3D-bioprinted constructs

To test the central premise of this study, 3DBP cell-laden structures need to exhibit sufficient stability to be cultured in vitro and be robust to withstand in vivo implantation. From a practical standpoint we chose a circular ring structure as the printing geometry, as the ring-shaped geometry is one of the most basic elements of many complex structures, such as hollow cylinder and sphere, and is also capable of enduring flexural and compressive stresses encountered by sub-dermal implants. Furthermore, since a ring structure cannot be stably realized using Ultrafoam due to forces exerted by the MSCs which will collapse the hollow center, this design paradigm also serves to demonstrate the unique features achievable by the proposed system.

Efficiency of output, expressed as the percentage of successful print of several objects of the same geometry in a sequence, is a critical parameter towards exploitation and clinical translation, especially when involving structures comprising of a high density of cells. Thus, we tested the utilization efficiency of the bioink by printing 1 mL and 0.5 mL of material using a ring with external diameter of 9 mm and a height of 2 mm as the model object. The output efficiency was 90% and 75% for 1 mL and 0.5 mL initial printing volume, respectively (**Fig. S2C, D**), indicating that CA-based bioinks offer high level of standardization, with both practical and economic advantages.

Since bone formation through EO proceeds via a cartilage template, both MSC-3DBP rings (volume 68.7 mm^3^) and MSC seeded at identical density on Ultrafoam (volume 84.8 mm^3^) were cultured in serum-free chondrogenic media supplemented with TGF-β3 for 3 weeks to induce the chondrogenic differentiation of MSC, followed by 2 weeks in medium designed to induce chondrocyte hypertrophy. Microcomputed tomography (µCT) images of both 3DBP rings and Ultrafoam discs showed that a mineral phase, still absent at 3 weeks, became evident at 5 weeks, i.e., after induction of hypertrophy (**Fig. 5A**) and was confirmed by Alizarin-Red staining (**Fig. 5B**). The pattern of matrix deposition in 3DBP samples was different and could be explained by their geometry which allows for diffusion of nutrition luminally and peripherally. 3DBP constructs exhibited key cartilaginous extracellular matrix (ECM) components, namely GAG (red in Safranin-O staining) and collagen type II (Col II, brown), throughout the structure, while in the Ultrafoam group ECM deposition was predominant in the outer regions with the formation of a necrotic core, likely due to the lack of access to nutrients and consistent with previous studies ^22^ **(Fig. 5B)**. Nevertheless, GAG/DNA values in Ultrafoam constructs were higher than the 3DBP constructs (**Fig. S4**), in spite of the more intense staining of 3DBP sections for GAG. Following exposure to hypertrophic media, 3DBP tissues expressed collagen type X (Col X, brown) a classical marker for hypertrophy and showed maturation of Col II and GAG deposition similar to what was observed using the Ultrafoam scaffold **(Fig. 5B)**. All the above findings support that MSCs can be differentiated into chondrocytes and subsequently undergo hypertrophy in CA-bioink derived 3DBP structures, thus forming the fundamental precursor tissue necessary for bone formation through EO.

**Fig. 5.**
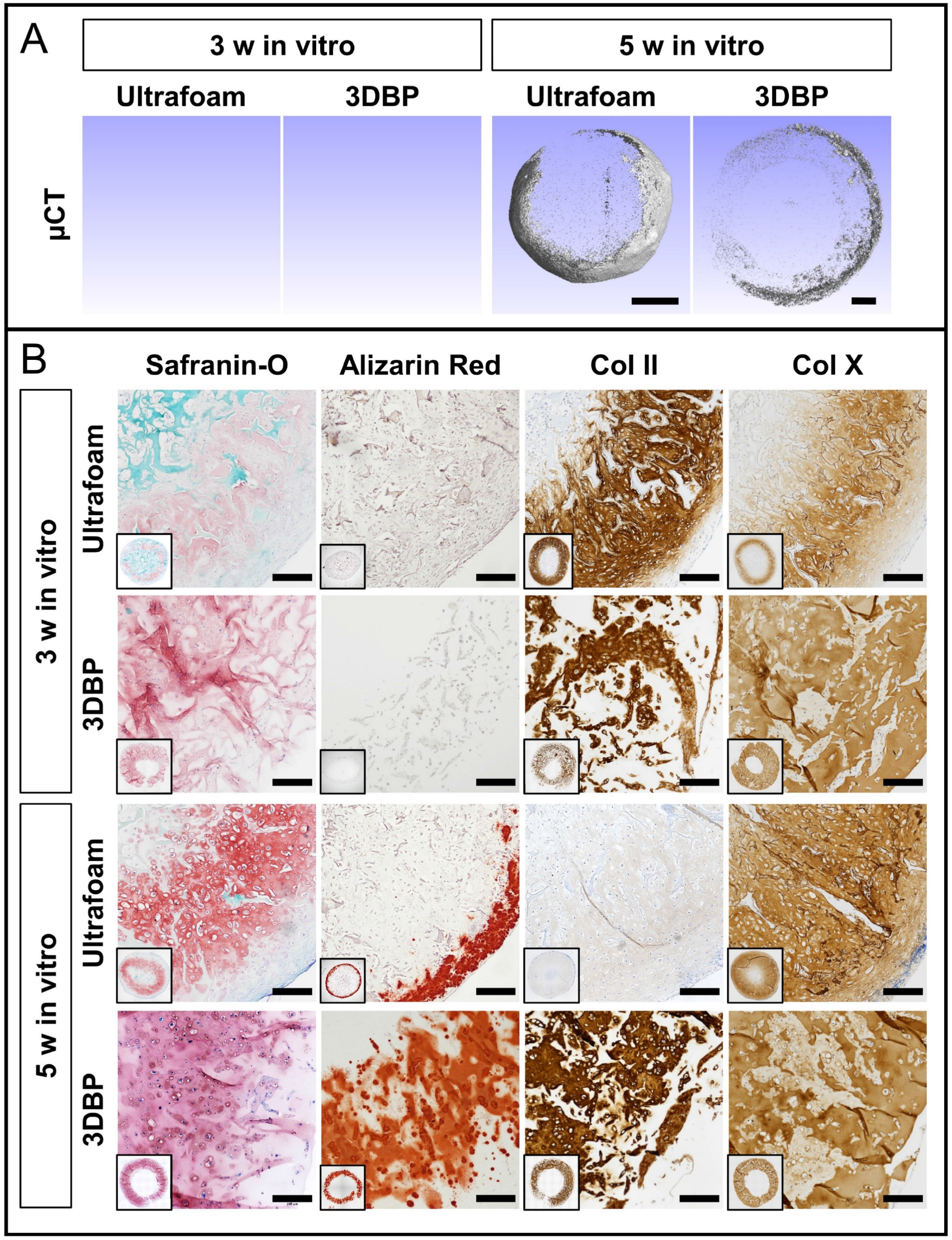
3DBP constructs undergo chondrogenesis and yield hypertrophic cartilage templates. (A) Three-dimensional reconstructed micro-computer tomographic (µCT) images of constructs after 3 weeks and 5 weeks of in vitro culture. Scale bars = 1mm. (B) Representative optical images of stained (Safranin-O, Alizarin-Red, Col I, and Col X) histological sections. Scale bars = 200 µm. (n = 3)

### Bioprinted hypertrophic cartilage remodels in vivo into bone structures of designed geometry

Based on the *in vitro* data, hypertrophic 3DBP and collagen constructs were implanted subcutaneously in nude mice to complete the EO program and harvested at 6 weeks and 12 weeks post-implantation. One important outcome was the stability of the 3DBP construct shape *in vivo*. Over the course of 12 weeks, all the 3DBP constructs remained intact. After 6 weeks, a gradual transformation of the constructs into osseous tissue was evident, with progressive maturation at 12 weeks. Interestingly, the osseous tissue deposition in the 3DBP constructs was annular (doughnut shaped), matching the printed structure, and was restricted to the prescribed original shape throughout the duration of *in vivo* maturation (**Fig. 6A**). After 12 weeks of *in vivo* remodeling, cortical bone was homogeneously distributed through the whole sample, similarly to the Ultrafoam group (**Fig. 6A**). The presence of bone tissue was confirmed by hematoxylin and eosin (HE) and Safranin-O staining, while immunohistochemistry for type II and type X collagen documented the resorption of cartilage tissue (**Fig. 6B**). The area positive for GAG (red area) had significantly shrunk after 6 weeks *in vivo* and was replaced by bone matrix (green area) in both 3DBP and Ultrafoam groups. Furthermore, a more pronounced staining for COL X in comparison to COL II was observed, indicating further maturation of the provisional cartilage matrix into the EO pathway. After 12 weeks of *in vivo* maturation, in addition to bone matrix (green counterstain in the Safranin-O staining) which was also confirmed by Masson’s trichrome staining (green area, **Fig. 7A**), marrow elements were evident (wine red part in Safranin-O staining) as well. Both osteoclasts and osteoblasts, which were assessed by staining for tartrate-resistant acid phosphatase (TRAP) and Osterix (OSX), respectively, were found throughout the samples (**Fig. 7A**). Direct contribution of the implanted MSCs in the bone formation was confirmed by positive staining of *in situ* hybridization for human Alu repeats, which are unique to human nuclei (**Fig. S5**). Based on quantitative analysis of µCT images, the average bone volume in 3DBP group increased more than 4-fold (from 2.5% at 6 weeks to 11% at 12 weeks), and in the Ultrafoam these values were 9.5% and 17.5%, respectively for not even 2-fold increase (**Fig. 7B**). Furthermore, it is quite remarkable that at 12-weeks the final volume of 3DBP sample was 3.6-fold greater than Ultrafoam sample (72 mm^3^ vs 20 mm^3^) (**Fig. 7C**), as the Ultrafoam sample had shrunk from the prescribed initial volume of 84 mm^3^ scaffold to 20 mm^3^, while the 3DBP samples maintained the prescribed initial volume (75 mm^3^ versus 68.7 mm^3^). In bone formation through EO, the remodeling of the provisional cartilage matrix is preceded by ingress of new blood vessels from existing surrounding vasculature. This step ensures the graft undergoes integration with the host tissue. CD31 staining in 6-week samples revealed the presence of CD31 positive structures in the exterior and inside of the lumen of the ring constructs and the margins of the controls that were in direct contact with the surrounding tissue. Such structures were identified in the center area of the constructs adjacent to regions that stained positive for osteocalcin, a major non-collagenous protein in bone matrix secreted by osteoblasts ^24^. At 12 weeks, staining of CD31 positive structures was more pronounced and found throughout the samples in both groups and were associated with host-derived cells indicating that both matrices are permissible to cell infiltration (**Fig. 8**).

**Fig. 6.**
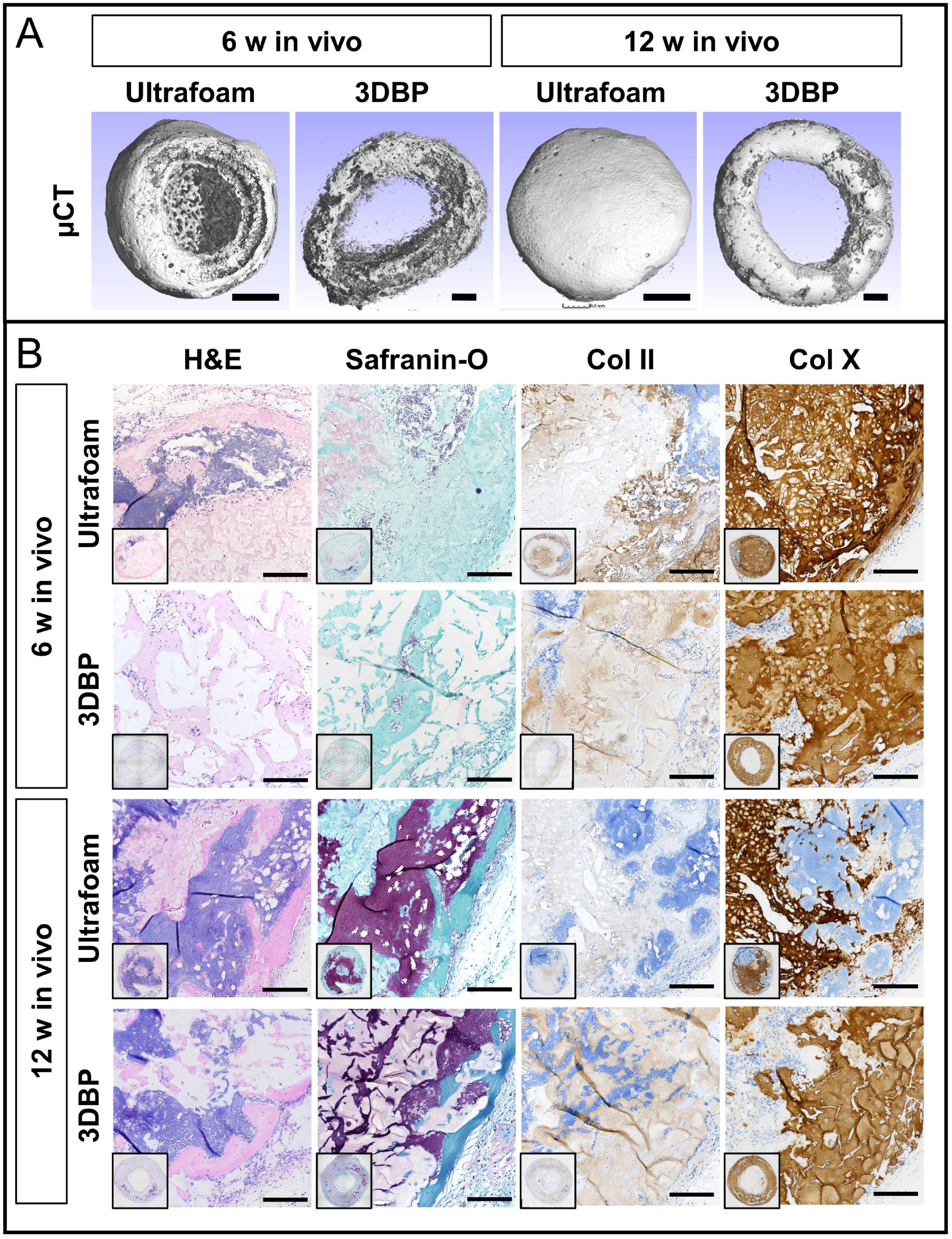
Implanted 3BBP hypertrophic cartilage templates remodel into bone of pre-prescribed shape through endochondral ossification in ectopic sites. (A) Three-dimensional reconstructed µCT images of constructs after ectopic implantation in nude mice for 6 weeks and 12 weeks. Scale bars = 1 mm. (B) Representative images of stained (H&E, Safranin-O, Col II, and Col X staining) histological sections. Scale bars = 200 µm. (n = 5 for Ultrafoam and 9 for 3DBP)

**Fig. 7.**
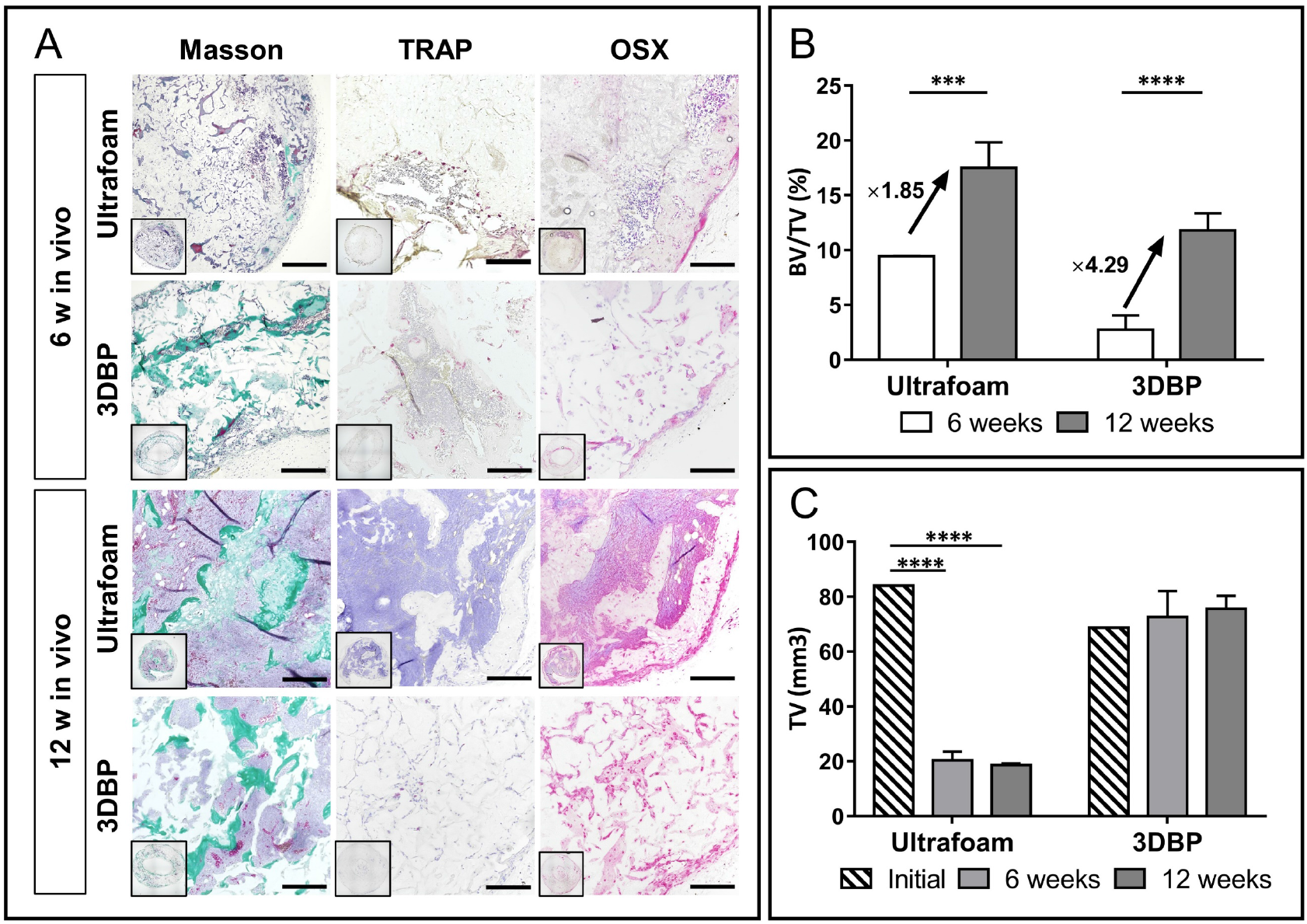
Remodeled 3DBP constructs retain the prescribed volume and possess key markers of bone. (A) Staining for bone-specific markers in explants after 6 and 12 weeks after in vivo maturation demonstrating the formation of mature bone in 3DBP constructs. Scale bars = 100 µm. (B) Proportion of bone volume to the total volume of the constructs indicates a superior fold-change in 3DBP group compared to Ultrafoam group. (C) The total volume of 3DBP constructs matched the designed structure and maintained unchanged through 12 weeks, while Ultrafoam collagen scaffolds shrunk 4 times compared to the initial prepared volume. (n = 5 for Ultrafoam and 9 for 3DBP, ***, *p*<0.001, ****, *p*<0.0001)

**Fig. 8.**
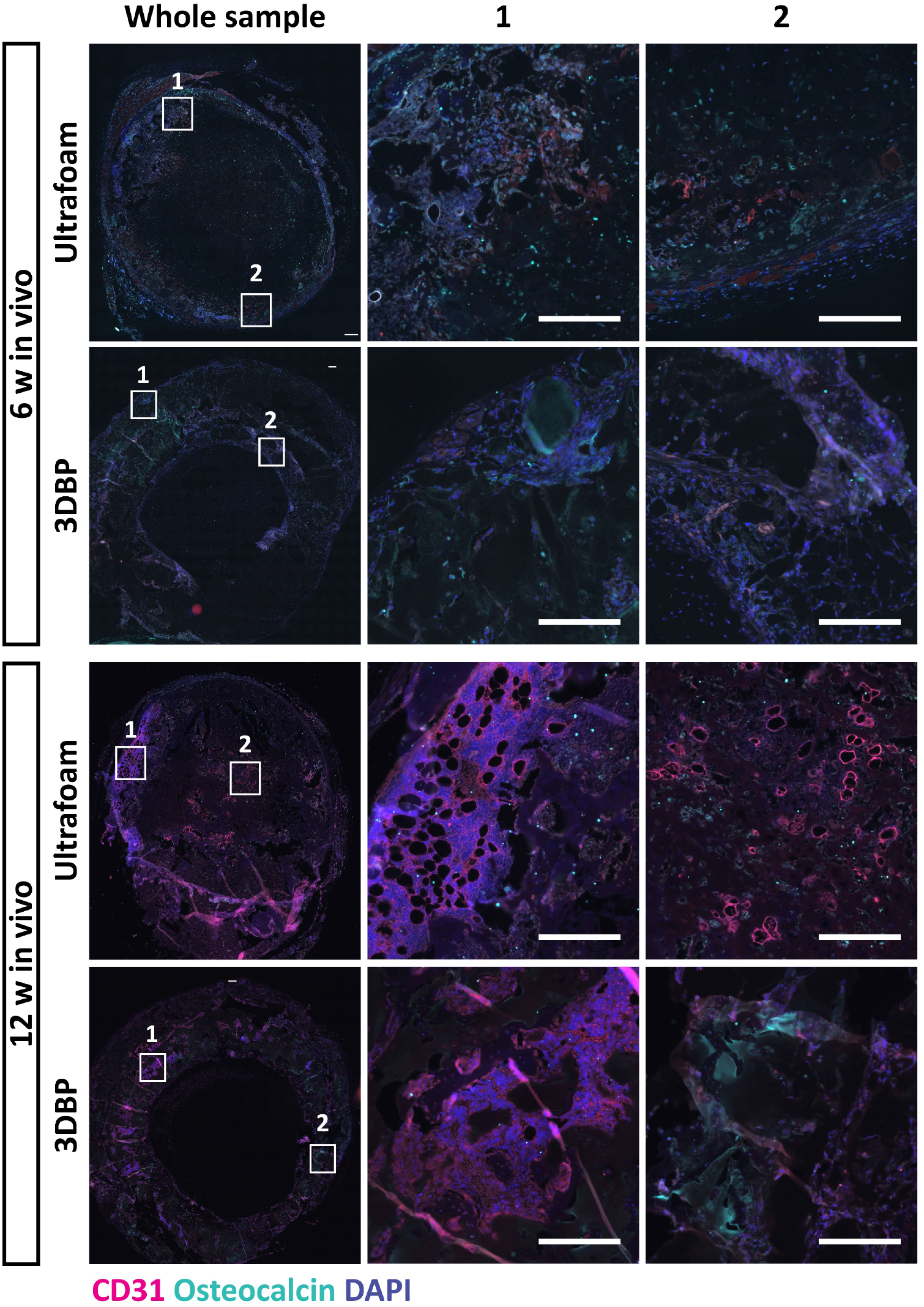
Bone tissue within 3DBP constructs is vascularized and show osteoblastic bone formation. After 6 weeks of implantation, osteocalcin, a marker associated with osteosynthesis was found interspersed in both Ultrafoam and 3DBP groups. Staining for CD31 a marker specific to endothelium associated with vasculature was observed predominantly in the edge areas of the implants that was directly in contact with the host environment (indicated by the white boxes). After 12 weeks, CD31 was found throughout the constructs associated with osteocalcin positive regions indicating the presence of vascularized bone. Scale bars = 200 µm.

## Discussion

In this study, we have demonstrated that 3D-printed hydrogel structures comprising of human MSCs can be successfully implanted and transformed into living bone tissue of pre-defined shape by combining 3D bioprinting with developmental tissue engineering. This was accomplished through the development of a CA-based bioink, that is capable of yielding single-matrix 3D-printed constructs of complex and anatomical shapes without the need for a support and fugitive phase and that can also incorporate a high cell density of cells (10’s of millions) while maintaining printability and post-printing stability. Formulating the bioink was a critical step in realizing the objectives of the study design. In our past effort we had identified onset of gelation close to physiological temperature, and a rapid gelation at low temperatures as two critical parameters for printing mechanically stable structures without support phases ^13^. The bioink formulation developed herein yielded structures that ranged from flat 2D objects to high aspect ratio twisted hollow cylinders, structures with defined high aspect ratio ridges (5:1 aspect ratio) and entirely solid objects. This range of complex geometries covers a significant landscape of the bone shape and thickness found in the human body, with the femoral cortical bone reaching thickness of up to 10 mm, and the flat bones of the face and skull ranging from 6 -10 mm in thickness ^25,26^. The phalange printed here as an example, is an 18-mm high tapering solid structure, with an elliptical base of 9 mm major axis and 6.5 mm minor axis corresponding to that of a human adults third finger middle phalange. This provides a solid platform to explore tissue engineering strategies targeting bone structures of pre-defined shapes and volumes.

EO is a process by which hypertrophic cartilage is remodeled in vivo into bone. Recapitulation of EO process through induction of hypertrophy in MSC-derived cartilage matrix, enabled the constructs to survive hypoxic conditions following implantation, with the printed structure providing the physical guidance for the formation of ossicles within a prescribed structure. Cartilaginous constructs for implantation were prepared by printing high density of MSCs (30 million cells per mL) and in vitro culturing under chondrogenic conditions followed by induction of hypertrophy. All 3DBP structures implanted in ectopic (subdermal) sites in mice survived the implantation intact over a 12-week period and underwent remodeling into bone. This was confirmed by staining for bone specific markers and the presence of invading vasculature localized around osteocalcin positive sites at 6 weeks and at 12 weeks.

However, several improvements have to be undertaken before 3D-bioprinted constructs can be viable for clinical use. In order to ensure bone formation throughout the construct, the bioink could be modified to possess hydrolysable linkages that can be proteolytically processed to promote cellular infiltration and tissue remodeling. Since vascularization is a pivotal step in EO process, engineering the bioink to present pro-angiogenic signals such as growth factors or incorporating genetically modified cells that secrete define soluble signals and decellularized ECM could provide a framework for optimization of the bioink performance in vivo. Recently, a devitalized tissue derived from human cell line of mesenchymal origin was shown to effectively and reproducibly form cartilage in vitro and showed accelerated remodeling into bone in vivo by promoting early recruitment of osteoprogenitors and osteoclasts ^27^. Finally, one also needs to consider donor specific changes in chondrogenic differential potential of MSCs and induction of senescence during expansion, both of which can impact ECM production ^28^.

In summary, the key findings of this study demonstrate that in principle 3DBP is a viable approach for fabricating living bone in predefined shape. From a clinical translation standpoint, process parameters and feasibility of embedding within a quality management system are paramount. In this regard, the single-matrix system presented here stands a great chance of clinical adaptability and meeting regulatory hurdles.

## Materials and Methods

### Synthesis of carboxylated agarose

Carboxylated agarose (CA) with 60% carboxylation was synthesized as previously described ^29^. Briefly, 10 g of native agarose (NA) type 1 (GeneOn, Germany) was transferred into a three-necked round bottom flask, equipped with a mechanical stirrer and pH meter. The reactor was heated up to 90 °C to dissolve the agarose and then cooled down to 0 °C in an ice bath under mechanical stirring to prevent the solution from gelling. The reactor was then charged with 300 mg TEMPO (Abcr, Germany), 1.5 g NaBr, and 37.5 mL NaOCl (15% v/v solution) under vigorous stirring. The pH of the solution was adjusted to pH 10.8 throughout the duration of the reaction, and the degree of carboxylation was controlled by the addition of predetermined volumes of NaOH solution (0.5 M). At the end of the reaction 1.5 g NaBH4 was added, and the solution was acidified to pH 8 and stirred for 1 h. The CA was precipitated by sequential addition of 150 g NaCl and 2 L ethanol, and the solid was collected by vacuum filtration and extracted using ethanol. Residual ethanol was removed by extensive dialysis against water and the CA was obtained as a white solid upon freeze-drying overnight. The degree of carboxylation was verified by the appearance of peaks associated with aliphatic carboxylic acid groups via FTIR (KBr) (νc, _C=O_ 1750 cm^−1^) (Bruker Optics, Germany) and NMR 300 MHz (13C: 180 ppm) (Bruker BioSpin, Germany).

### Rheological tests and compression tests

All the rheological tests were performed using a Kinexus-PRO+ rotary rheometer (Malvern, United Kingdom). A 4° cone plate with 40 mm in diameter and a plate were used in examinations of viscosity and shear stress. Except for temperature sweeping tests, the samples were loaded at a different temperature, samples for the other tests were loaded at 45 °C, and then maintained at 37 °C to reach equilibrium before tests. In frequency sweeping tests, oscillation with 1% strain ranging from 0.1 Hz to 10 Hz was exerted on samples. In shear stress ramping tests, sheared stress was controlled from 0 to 200 Pa. In time sweeping tests, samples were oscillated with 1% shear strain at 1 Hz for 30 minutes. In temperature ramping-down tests, samples were first loaded to lower plate at 45 °C and maintained for 5 minutes to reach equilibrium before being further cooled down to 4 °C at the rate of 2 °C /min, and in temperature ramping-up tests, samples were first loaded to lower plate at 37 °C and then maintained for 5 minutes before further heated up to 37 during tests. For temperature sweeping tests, oscillation was exerted with 1% shear strain at 0.1 Hz.

The unconfined compression tests were performed by using a Kinexus-PRO+ rotary rheometer (Malvern, United Kingdom). The bioink was loaded on the lower plate at 37 °C. And the upper parallel plate with a radius of 20 mm went down to initial height. Samples were trimmed to fit the diameter of the upper plate before tests, and then cooled down to 4 °C to form hydrogel. For compressing test, samples were first brought to the testing temperature and kept for 5 minutes for equilibrium. Then the upper plate went down at the speed of 0.001 mm/s to a given height to compress the hydrogel. The upper plate stopped either at the given gap or when the detected normal force reached 50 N. The distance between the upper plate and lower plate and normal force were recorded during the tests. The compressive modulus was calculated by using formula S1 (Supplementary materials) based on the data in the linear viscoelastic region of the hydrogel.

### Mesenchymal stromal cell isolation and in vitro culture

Human bone marrow-derived mesenchymal stromal cells (MSCs) were isolated from bone marrow aspirates of healthy donors. The tissue samples were obtained during orthopedic procedures under the general informed consent of the University Hospital Basel and in accordance with the local ethical committee’s regulations. MSCs were expanded with α-minimum essential medium (α-MEM)-based media (Gibco, Germany) containing 10% FBS (Gibco, Germany), 100 mM HEPES (PAN-Biotech, Germany), 100 U/mL penicillin, 100 mg/mL streptomycin (PAN-Biotech, Germany), and 5 ng/mL FGF-2 (R&D systems) (MSC expansion media).

### Lentiviral transduction of MSCs

Lentiviral particles containing shRNA constructs (pLVX-tdTomato) was produced in HEK293 cells, by co-transfecting lentiviral vector and packaging vectors using polyethyleneimine (Mw 25,000 Sigma, Germany) as the transfection reagent. For transfection, 30 µg of DNA (4:3:1 of transfer vector, packaging coding vector (pCMVdR8.74) and envelope coding vector (pMD2.G)) was diluted in 250 µL Opti-MEM (Invitrogen, Germany) and 11.25 µL of polyethyleneimine (1 mg/mL) was added to the solution, and the resulting mixture was incubated for 25 min at room temperature prior to adding to HEK293 cells. The medium was changed after 16 h to MSC expansion medium, and 64 h after transfection, the viral supernatants were collected and filtered through a sterile 0.22-μm syringe filter (Millipore, Germany). Viral particles were then added to target cells (MSC) and after three days of transduction infected cells were selected by flow cytometry sorting for tdTomato vector.

### CA-based Bioink preparation and 3D culture of MSCs

Lyophilized CA and NA powder were weighed in a syringe with thread at concentrations of 9.5% (w/v) and 0.5% (w/v) respectively. Then, Dulbecco’s phosphate-buffered saline (DPBS) was added to the syringe to dissolve the solute mixture at 95 °C water bath until a transparent solution. The CA/NA solution was then kept in 45 °C for 10 minutes before use. MSCs were digested with 0.05% trypsin /0.02% EDTA for 5 minutes and then neutralized with DMEM containing 10% FBS. Suspension of MSCs was then centrifuged at 800 rpm for 5 minutes, and the supernatant was removed. The pellet of MSCs was then dispersed and resuspended in MSC-expansion media as above-mentioned. For 3D seeding, tdTomato-MSCs with concentrations at 10 × 10^6^ cells/mL, 30 × 10^6^ cells/mL, and 60 × 10^6^ cells/mL were used. The cell-laden bioinks were then kept in 42 °C before transferred into 96-well plates. The 96-well plate was then kept in fridge for 2 minutes for gelation. After that media was added on top of hydrogels, and the cell-laden hydrogels were cultured at 37 °C in cell incubator supplied with 5% CO_2_.

### 3DBP of MSCs

CA-based bioink and suspension of MSCs were prepared as previously described. For mixing CA-based bioink and MSCs, MSCs suspension at a concentration of 30 × 10^6^ cells/mL or 60 × 10^6^ cells/mL was mixed with CA-based bioink for *in vitro* differentiation. The cell-laden bioink was transferred to the printer cartridge before printing. Printing procedure was carried out using an in-house modified Inkredible+ 3D bioprinter (Cellink, Stockholm, Sweden) with modifications as previously described ^13^. The temperature of cartridge and nozzle was set as 37 °C, while temperature of the print bed was set as 4 °C, throughout the printing. 3D model of mandibular bone was reconstructed from a patients’ CT scan, and the model of vertebral body and finger phalange were downloaded online licensed under Creative Commons ^30,31^. Ring-shaped constructs with 9 mm in outer diameter, 7.2 mm in inner diameter, and 3 mm in height (68.7 mm^3^) were printed and then equilibrated for 15-second on the print bed to ensure physical crosslinking, before being transferred to 12-well plate with agarose-coated bottom, following which 2 mL culture media was added and then cultured in appropriate media. Ultrafoam™ collagen scaffolds (6-mm diameter, 3mm high (84.8 mm^3^) seeded with MSCs a density of 30 × 10^6^ /mm^3^ served as control.

### Live/dead assay

Bioink was mixed with MSCs at a final concentration of 1 × 10^6^ cells/mL and printed. The printed constructs were washed twice with DPBS before culture in MSC expansion media. For cell viability determination, samples were collected at certain time points and stained with 0.5 µL/mL Calcein-AM (Life Technologies, Germany) and 2 µL/mL ethidium homodimer-1 (Life Technologies, Germany) in DPBS, and incubated for 30 min in cell incubator before instant imaging. The samples were integrally scanned on one layer by fluorescent imaging capture (Zeiss Observer Z1, Carl Zeiss, Germany). Then four random areas were scanned layer-by-layer at an interval at 60 µm. The counting procedure was performed in ImageJ software (National Institutes of Health, USA).

### Quantification of MSC aggregates and aggregate area

The number of singles cells and clusters in a field view were manually counted to establish a numerical value for aggregation in percent. The area of single MSCs and MSC aggregates was computed from fluorescence images by drawing an outline of the object and using the area function in Zeiss Zen Blue software.

### Differentiation of hypertrophic cartilaginous templates

Differentiation of MSCs into hypertrophic chondrocytes was carried out by incubation in successive chondrogenic and hypertrophic media for 3 weeks and 2 weeks respectively with two media changes every week. Both are serum-free media containing high glucose DMEM supplemented with 1mM sodium pyruvate (Gibco™), 10mM HEPES buffer (Gibco™), penicillin-streptomycin-L-Glutamine (100X, Gibco™), Insulin-Transferrin-Selenium-A (Gibco™) and Human Serum Albumin 0.12% (CSL Behring). Chondrogenic medium is supplemented with 0.1 mM ascorbic acid (A5960, Sigma), 10^−7^ M dexamethasone (D4902, Sigma) and 10 ng/mL TGF-β3 (Novartis). Hypertrophic medium is supplemented with 50 nM L-thyroxine, 10mM β-glycerophosphate (Sigma), 10^−8^M dexamethasone, 0.1mM ascorbic acid and 50 pg/mL IL-1β (Sigma).

### In vivo implantation

All animal studies were approved by the Swiss Federal Veterinary Office (permit 1797). Analgesia was provided 1h prior to surgery by subcutaneous injection of Buprenorphine (0.1 mg/kg body weight). Anesthesia with isoflurane (2.5%) was maintained on demand with oxygen as a carrier (0,6 L/min). Six to ten weeks old female CD-1 nude mice were used, two midline incisions (c.a. 5 mm) of the dorsal skin were performed using scissors under sterile conditions and up to four tissues were implanted per animal. Wound closing was done using sterile clips which were removed 7 to 10 days post-surgery. Animals were sacrificed using CO2 inhalation followed by cervical dislocation before tissue retrieval.

### Micro Computed Tomography

Samples were fixed using Formalin 4% overnight and immobilized in histogel for analysis. Microtomography of the explants was performed using a tungsten x-ray source at 70 kV and 260 µA with an aluminum filter of 0.5 mm (Nanotome, GE, USA). Transmission images were acquired for 360° with an incremental step size of 0.25°. Volumes were reconstructed using a modified Feldkamp algorithm (software supplied by manufacturer) at a voxel size of 10 µm. Thresholding, segmentation and 3D measurements were performed using the VG Studio Max software. After microtomography, samples were decalcified in 15% EDTA solution (Sigma Aldrich) before histology.

### Histological characterization

Samples were embedded in paraffin and 5 µm thick sections were prepared using a microtome (Microm, HM430, Thermo Scientific). Safranin-O, Alizarin red, hematoxylin/eosin, Masson Trichrome, TRAP, Osterix and Alu stainings were performed as previously described (Scotti et al., PNAS 2013). Collagen type II (Reactivity: human, MP Biomedicals, 63171) and X (Reactivity: human, Invitrogen 14-9771-80) stainings were performed with Ventana Discovery Ultra (Roche Diagnostics (Suisse) SA) automated slide stainer. In brief, tissue sections were deparaffinized and rehydrated. Antigens were retrieved by a protease (Protease 3, ref. 760-2020, Ventana) digestion for 20 to 44 minutes at 37 °C. Primary antibody was manually applied and incubated for 1 hour at 37 °C. After washing, the secondary antibody was incubated for 1 hour at 37 °C. Detection step was performed with the Ventana DISCOVERY ChromoMap DAB (ref. 760-159 Ventana) detection kit. Afterwards, the slides were counterstained with hematoxylin II, followed by the bluing reagent (respectively ref. 790-2208 and 760-2037, Ventana). Sections were then dehydrated, cleared and mounted with permanent mounting and coverslips.

### Immunofluorescence staining of CD31 and OCN

CD31 and Osteocalcin (OCN) staining was performed on paraffin slides following overnight paraffin melting at 60°C, deparaffinization/rehydration bathes, and antigen retrieval at 98°C for 15 minutes (KOS microwave tissue processor, Milestone) in citrate buffer pH6.0 (Sigma, C999). Prior to staining, tissues where permeabilized using 0.3% Triton X-100 (Sigma, 93443) in PBS and blocked for unspecific binding using 1% Bovine Serum Albumin (BSA) (Sigma A3803) + 5% goat serum (Invitrogen, 16210-064) in PBS + 0,02% Tween-20 (PBS-T) (Sigma, P-1379). Tissues were then stained with mouse anti-CD31 antibody (Reactivity: human, Abcam, ab9498) at a dilution 1:2000 and rabbit anti-OCN antibody (Reactivity: human, Invitrogen, PA5-96529) at a dilution 1:100 in 1% BSA in PBS-T for 1h00 at room temperature. Secondary antibodies goat anti-mouse Alexa 555 (Life Technologies, A21422) and goat anti-rabbit Alexa 647 (Life Technologies, A21244) at a concentration 1:200 in 1% BSA in PBS-T were incubated for 45 min at room temperature. DAPI was applied for 5 min at the end of the staining. Washing steps were performed with PBS and slides were mounted using Dako Faramount Aqueous mounting medium Ready to use (S3025).

### Statistical analysis

All values are reported as mean ± standard deviation. Statistical analysis was carried out using Origin2019b, and two-way ANOVA was used to compare the difference between each two sets of data. A *p*-value of ≤ 0.05 was considered statistically significant.

## Acknowledgments

The authors wish to thank MAPTECH HOLDINGS UG for generous access to the Kinexus Pro+ rotary rheometer, the DBM Histology Core Facility, Universitätsspital Basel for the support with staining using Ventana, and Dr. Florian Thieringer from Mund-, Kiefer-und Gesichtschirurgie, Universitätsspital Basel for providing the reconstructed 3D jaw model from CT scans.

## Funding

The excellence initiative of the German Federal and State governments (EXC 294) China Scholarship Council Doctoral Fellowship (Y.G.) The Swiss National Science Foundation (Grant No. 310030-133110 to I.M.)

## Author contributions

Conceptualization: PS, IM, AB, YG, SP

Methodology: YG, SP, LA, MS, PS, AB, IM

Investigation: YG, SP, FTR, LA

Resources: PS, IM Visualization: YG, SP, PS

Supervision: PS, AB, IM

Writing—original draft: YG, PS, SP

Writing—review & editing: YG, SP, FTR, LA, MS, AB, IM, PS

## Competing interests

PS is an inventor on several patents on the use of carboxylated agarose hydrogels. All other authors declare they have no competing interests.

## Notes

### Competing Interest Statement

Prasad Shastri is an inventor on several patents on the use of carboxylated agarose hydrogels. All other authors declare they have no competing interests.

