## Supplementary Information for "Towards 3D-bioprinting of osseous tissue of pre-defined shape using single-matrix cell-bioink constructs"

**This PDF file includes:**

Supplementary Text
Figs. S1 to S5

### Supplementary Text

Materials and methods

#### Mechanical properties

The compressive modulus is calculated based on the following formula,

$$E=(\frac{\Delta F}{\Delta L})\frac{L_0}{A_r} \qquad \text{(Formula S1)}$$

as in the equation $L_0$ denotes the initial height of the hydrogel, and $A_r$ denotes the surface area of the applied plate of during the tests, with $\Delta F$ and $\Delta L$ as the values corresponding to the changes measured between two specific points in linear viscoelastic region of the hydrogels.

#### Glycosaminoglycans (GAG) measurements

Samples were digested overnight at 56 °C in 1 ml of proteinase K solution (Sigma Aldrich, P2308), and 100 µl of the resulting digested solution was incubated with 1 ml of DMMB solution (16 mg/l dimethyl-methylene blue, 6 mM sodium formate, 200 mM GuHCL, all from Sigma Aldrich, pH 3.0) on a shaker at room temperature for 30 minutes. After centrifugation, precipitated DMMB-GAG complexes were dissolved in decomplexaion solution (4 M GuHCL, 50 mM Na-Acetate, 10% Propan-1-ol, all from Sigma Aldrich, pH 6.8) at 60 °C for 15 minutes. Absorption was measured at 656 nm and corresponding GAG concentrations were calculated using a standard curve prepared with purified bovine chondroitin sulfate (Sigma Aldrich).

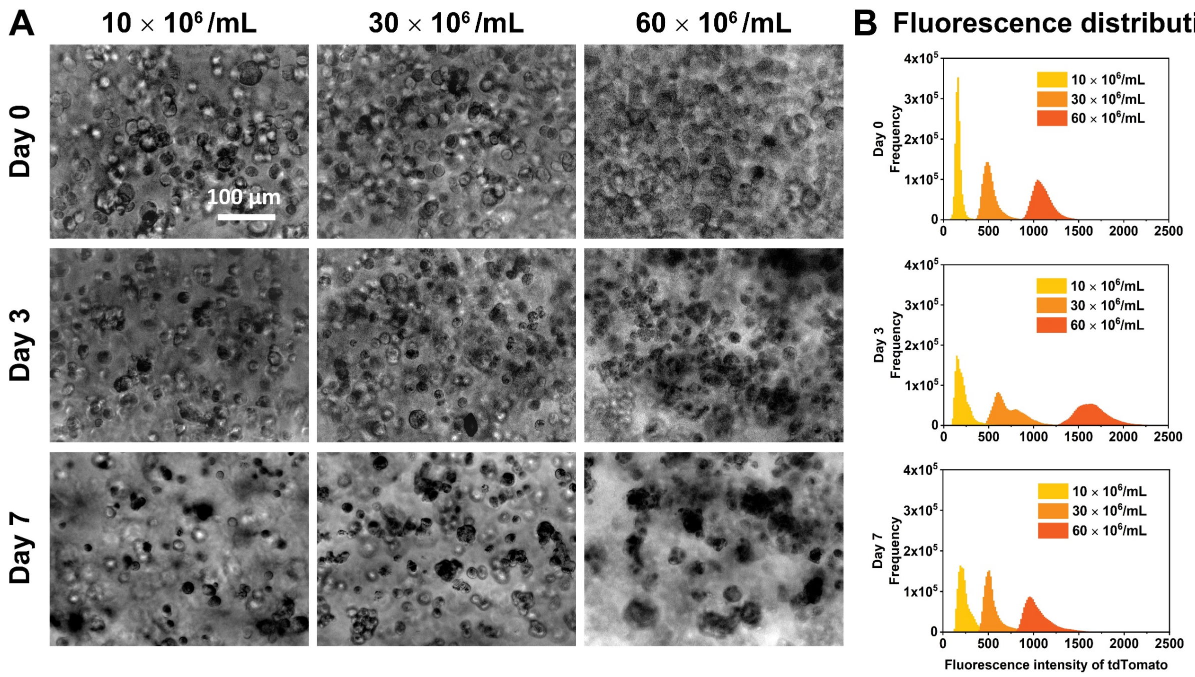


**Fig. S1.**

(A) Morphology presented under bright field of BM-MSCs inside CA bioink with different seeding cell density from day 0 to day 7. Cells presented inflated round morphology in all cell densities instantly after seeding (Day 0), and in 60 × $10^6$/mL group, more cells were found adjacent to each other. After 3 days of seeding, all cells shrank to smaller round shape, and more cell aggregates could be found with increasing cell density. (B) Intensity of fluorescence from MSCs distributed in bioink on from seeding day till $7^{th}$ day after seeding.

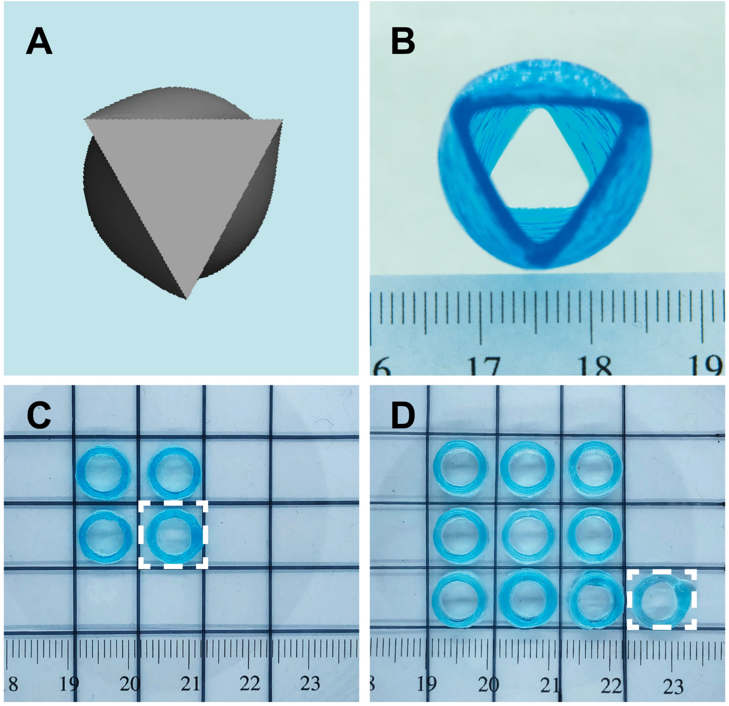


**Fig. S2.**

The top view of an irregular tube with two triangular openings rotated by 60° with respect each other, presenting a star-of-David shape configuration (A, B). Printing efficiency with 0.5 mL bioink (C) and 1 mL bioink (D). The printed model is a ring with dimensions as follows: 9 mm in outer diameter, 7.2 mm in inner diameter, and 3 mm in height. For 0.5 mL bioink, 3 complete tubes and 1 incomplete tube (in rectangle frame with white dotted line in A) were printed, and for 1 mL bioink, 9 complete rings, and 1 incomplete ring were printed (in rectangle frame with white dotted line in B). All the rings were printed one by another without any removed samples.

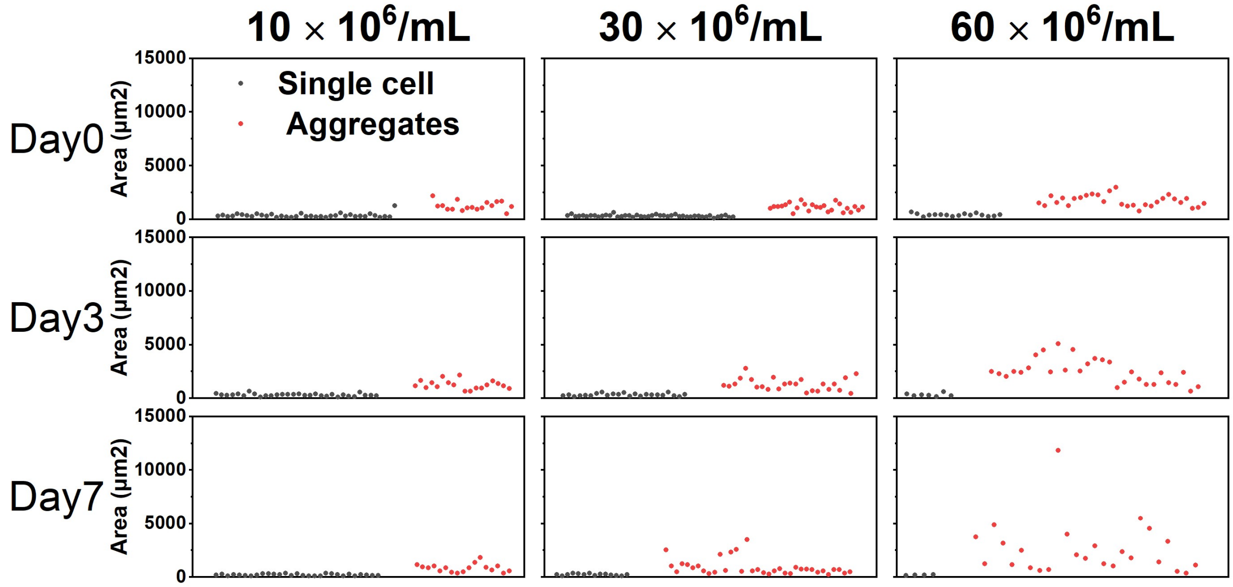


**Fig. S3.**

Distribution of single cells and cell aggregates inside CA bioink with different seeding cell density with time course. The average size of cell aggregates was found to increase in 30 × $10^6$/mL, and 60 × $10^6$/mL groups from day 0 to day 7 after seeding, and the size of aggregates differed more 60 × $10^6$/mL group since day 3. The average size of cell aggregates showed slight change during culture.

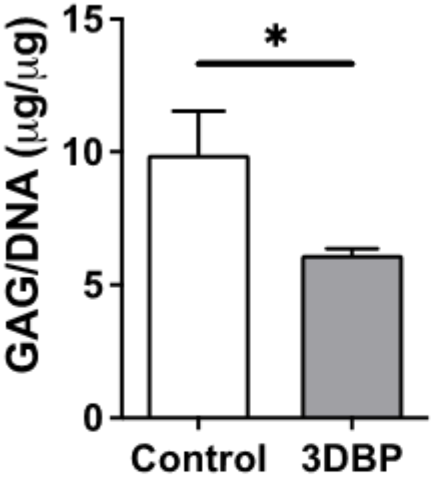


**Fig. S4.**

GAG/DNA analysis after 5 weeks of in vitro culture. The levels of GAG per DNA were 9.91 μg/ μg, and 6.13 μg/ μg for control and 3DBP, respectively. (n=2 for control, and n=5 for 3DBP)

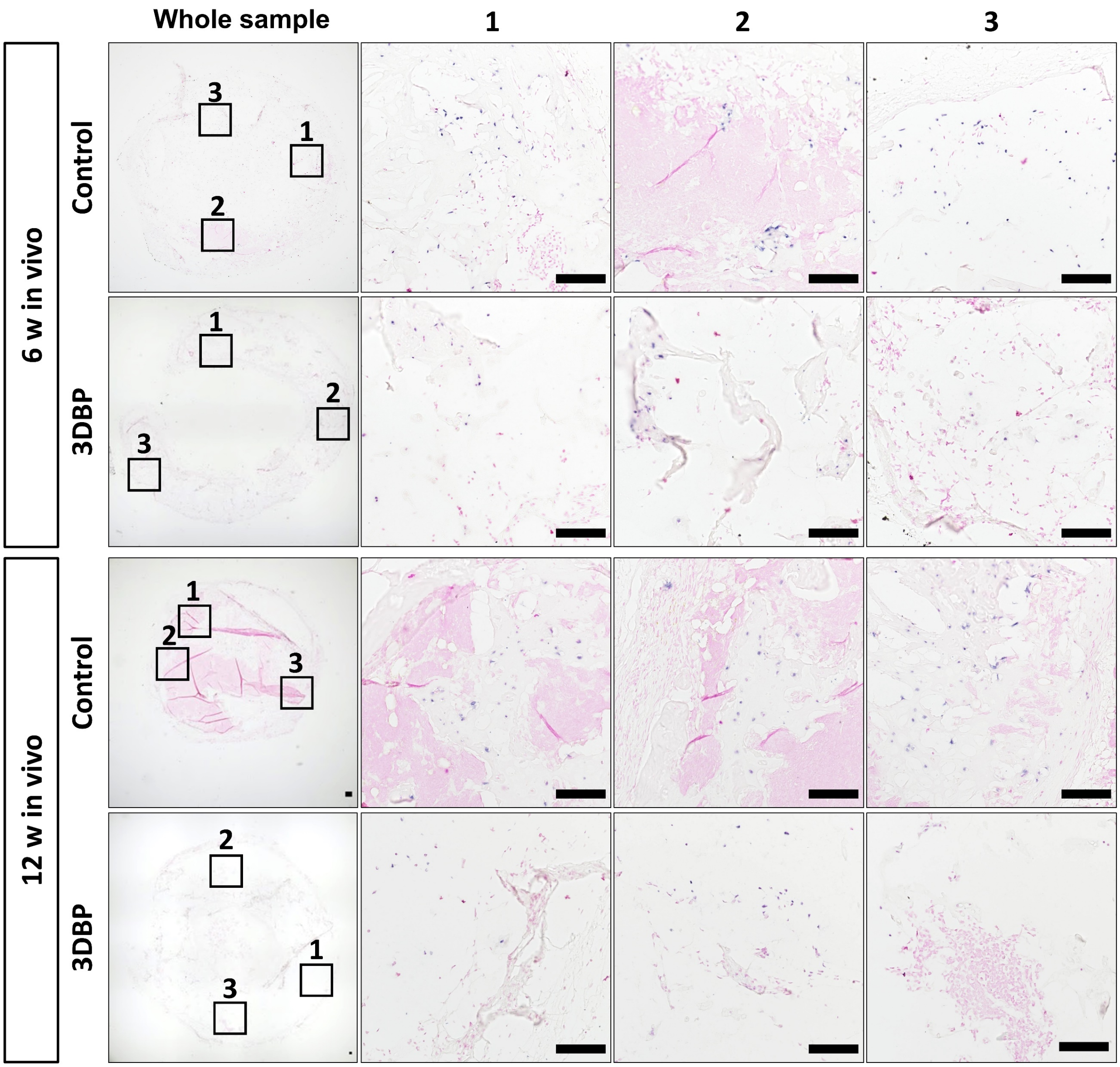


**Fig. S5.**

In situ hybridization for human Alu repeats in samples retrieved after 6-week and 12-week implantation. Scale bar = 100 µm.
